# Structural and Functional Characterization of Encapsulin-Targeted Double Ferritin Fold Ferroxidases

**DOI:** 10.1101/2025.09.13.676036

**Authors:** Arina Anuchina, Alina Remeeva, Ilia Natarov, Anna Yudenko, Rahaf Al Ebrahim, Pavel Shishkin, Vsevolod Sudarev, Valeriia Matveeva, Oleg Semenov, Elizaveta Kuznetsova, Andrey Nikolaev, Ivan Bezruchko, Daria Kuklina, Elizaveta Dronova, Na Li, Yury Ryzhykau, Nikolai N. Sluchanko, Yuqi Yang, Valentin Borshchevskiy, Alexey Vlasov, Sergey Bazhenov, Ilya Manukhov, Ivan Gushchin

## Abstract

Ferritins are a widespread family of proteins involved in iron homeostasis. While classic ferritins consist of four α-helices and form 24-meric nanocages, related ferritin-like proteins display other types of assemblies and sometimes lack any iron storage capacity. Here, by analyzing the available genomic data, we identify a family of double ferritin-like proteins (DFLPs) composed of two four-helical domains, which arose by duplication of a ferritin fold protein. We characterize representative DFLPs from *Thermocrinis minervae* and *Caldanaerovirga acetigignens*, TmDFLP and CaDFLP, and show that they form homodimers and bind heme. We determine the X-ray structure of TmDFLP and demonstrate its ferroxidase activity. Furthermore, we show that some DFLPs, including TmDFLP and CaDFLP, are highly likely to be targeted into encapsulin shells. Our work expands the range of known iron metabolism systems and highlights the power of genome mining for discovery of new proteins.

**Graphical Abstract for Table of Contents:** 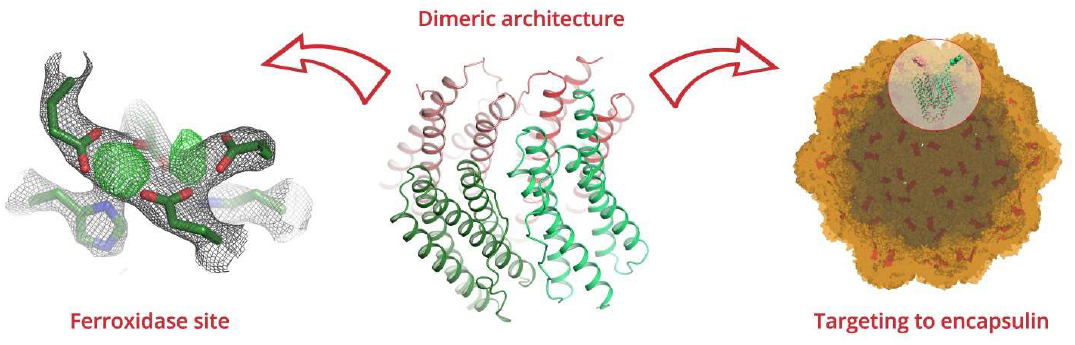

A family of double ferritin-like proteins (DFLPs) composed of two four-helical domains is described and investigated. DFLPs are shown to form homodimers, bind heme, possess diiron sites and display ferroxidase activity. Some DFLPs are shown to be targeted to encapsulin shells as core or secondary cargo, thus representing a new type of iron metabolism systems.

## Introduction

Ferritins are a large group of proteins widespread among all living organisms from Archaea to higher eukaryotes^[1–3]^. They participate in iron homeostasis by oxidizing Fe^2+^ ions (thus being ferroxidases), storing the resulting Fe^3+^ ions, and releasing them back when needed. Each protomer is organized as a four-helical bundle, which is also characteristic for a superfamily of ferritin-like proteins (FLPs) that includes rubrerythrins, manganese catalases, fatty acid desaturases, several types of oxygenases, R2-like ligand binding oxidases and ribonucleotide reductases^[3,4]^. The iron oxidation reaction proceeds at the conserved diiron site^[5]^.

Ferritins are divided into three subfamilies which differ by functions and multimer structures. Bacterioferritins (BFRs) are prokaryotic ferritins, which form 24-meric nanocages and have one heme B group per two ferritin monomers^[1,6]^. DNA binding protein from starved cells (Dps) forms 12-mers, has the smallest cage size among ferritins and contains no heme. In addition to the role in iron metabolism, Dps is involved in reorganization of DNA during stress conditions^[7–9]^. The third family, classic ferritins (FTNs), comprises 24-meric ferritins that do not bind heme and are found in bacteria (sometimes designated as bacterial non-heme ferritins) and eukaryotes^[3,10]^. In particular, higher vertebrate ferritins are typically heteromeric, containing two types of subunits: heavy and light chain^[2,11]^. Prokaryotic ferritins and ferritins from lower eukaryotes were considered to be homooligomeric until recently. However, recent studies demonstrate that 24-mer ferritin cages from *Pseudomonas aeruginosa* and *Acinetobacter baumannii* are heterooligomeric and consist of bacterioferritin and ferritin subunits. In comparison to Dps, ferritins and bacterioferritins function mainly as iron storage cages^[12–14]^.

Besides ferritins, a number of related ferroxidases were identified previously. These proteins form various oligomers (dimers to decamers), but do not form nanocages themselves and are embedded into encapsulin shells^[15,16]^. While some of these ferroxidases have been characterized functionally and structurally^[17–20]^, others remain understudied. Here, we identify a group of unique FLPs that contain a duplicated four-helical ferritin-like domain, which we designate double ferritin-like proteins (DFLPs). We present a combined bioinformatic and experimental study, which demonstrates that DFLPs are dimeric heme-binding encapsulin-targeted ferroxidases and thus represent a new bacterial iron metabolism system.

## Results and Discussion

### Identification and phylogenetic analysis of DFLPs

We began our study by analyzing the ferritin family sequences. At the moment of the analysis, initiated on January 25th, 2024, 85049 proteins were annotated as having the IPR008331 domain (“Ferritin/DPS protein domain”) in the InterPro database^[21]^. Similarly to previous studies^[3,2,17]^, our phylogenetic analysis reveals three major clades - a clade with Dps-like proteins, a clade with diverse proteins related to bacterioferritins, and closely related clades of eukaryotic ferritins and bacterial non-heme ferritins (Figure 1).

**Figure 1.**
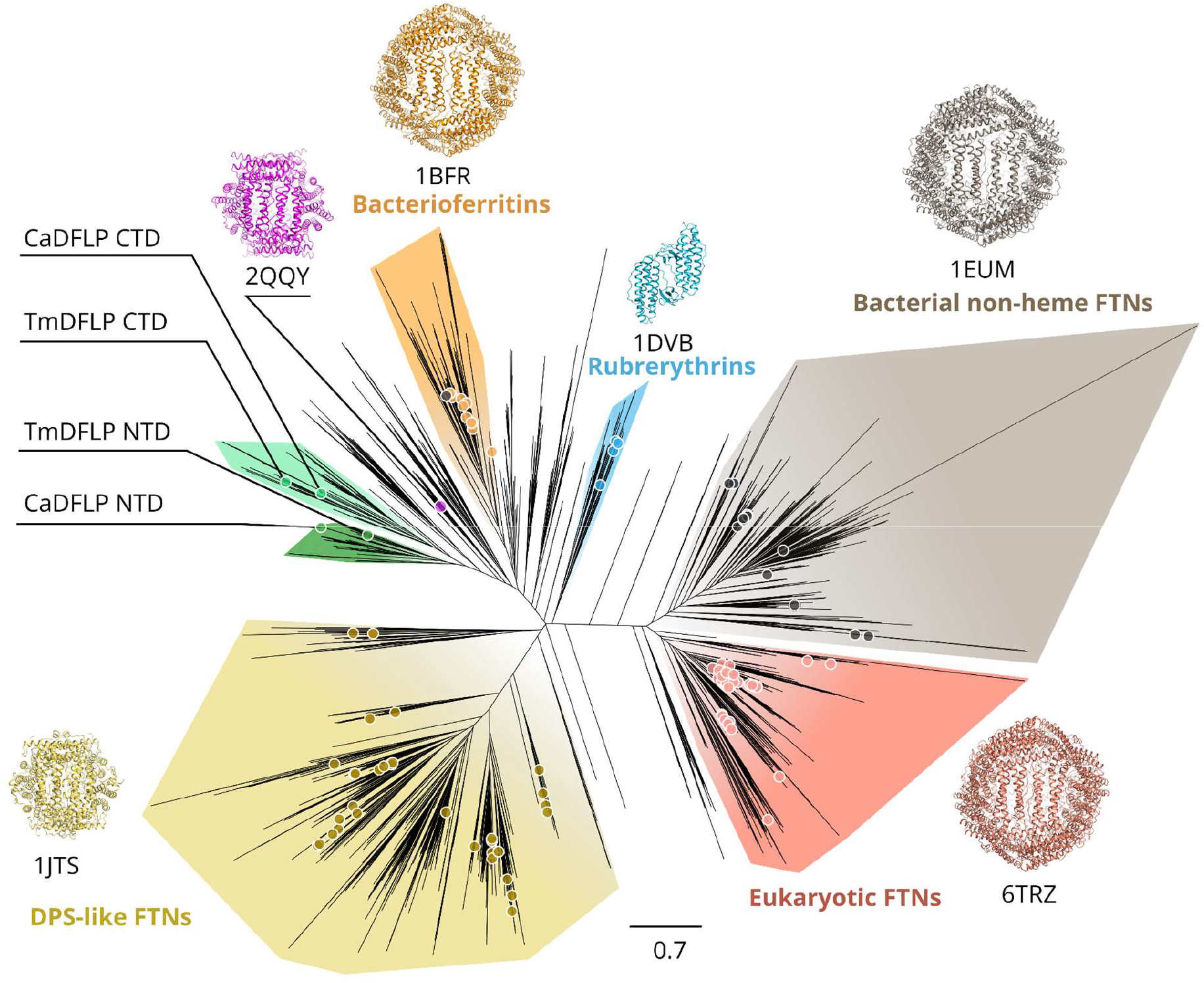
Phylogenetic tree of ferritin-like proteins. Structurally characterized proteins and DFLPs characterized in this work are marked with circles. N- and C-terminal domains of DFLPs form distinct clades marked in dark green and light green, respectively. The closest structurally characterized homolog of DFLPs is a 12-meric ferritin-like protein from *Bacillus anthracis* of unknown function (PDB ID 2QQY^[22]^). Rubrerythrins, which are not considered as members of the ferritin family, are added as an outgroup.

Among the analyzed sequences, we detected 157 double ferritin-like proteins (DFLPs), composed of two four-helical ferritin domains. We analyzed N-terminal domains (NTDs) and C-terminal domains (CTDs) of DFLPs separately and observed that NTDs and CTDs form distinct clusters having the same origin in the overall phylogenetic tree, close to the previously characterized bacterioferritin clade (Figure 1). Thus, DFLPs most likely arose due to duplication of a bacterioferritin-like gene and then underwent further speciation in different organisms. For a more detailed investigation of the features of DFLP clade proteins, we selected two proteins from thermophilic bacteria, *Caldanaerovirga acetigignens* (CaDFLP) and *Thermocrinis minervae* (TmDFLP), because proteins from thermophiles often display high stability, which is beneficial for experimental studies^[23]^. For both *C. acetigignens* and *T. minervae*, DFLPs are the only ferritin domain-containing proteins found in the genome.

### Binding of heme by DFLPs

We expressed CaDFLP and TmDFLP in *Escherichia coli* and observed slight pink coloring of the purified samples, with the absorption spectra revealing characteristic heme features, more pronounced for TmDFLP (Figure 2A). Position of the Soret peak varied from 412 nm for an oxidized sample to 430 nm for a DTT-reduced sample, as expected for protein-bound heme^[24]^ (Figure 2B). Based on the absorption values at 280 and 410-430 nm, we estimate the heme:protein molar ratios as ∼3% and ∼8% for CaDFLP and TmDFLP, respectively. Such sub-stoichiometric binding might be a consequence of suboptimal expression conditions, heme loss during purification, or selectivity of DFLPs to other heme types not present in *E. coli*.

**Figure 2.**
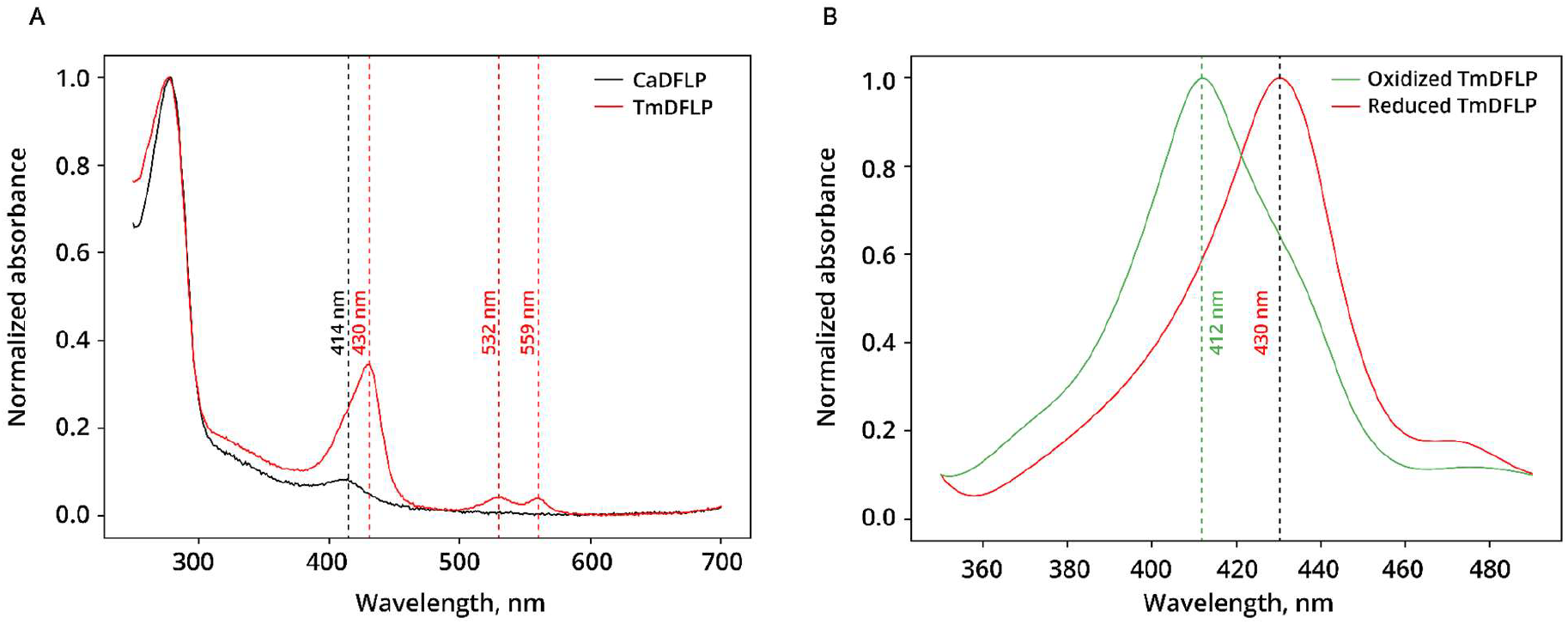
Absorbance spectra of CaDFLP and TmDFLP expressed in *E. coli*. A) UV-Vis absorption spectra of CaDFLP and TmDFLP following metal-affinity purification. B) Dependence of the TmDFLP Soret peak position on redox conditions.

### Oligomeric state of DFLPs

Unexpectedly, we observed that during purification TmDFLP and CaDFLP elute not as high molecular weight nanocages, but rather as relatively low molecular weight species. Multi-angle light scattering coupled with size-exclusion chromatography (SEC-MALS) indicates that CaDFLP elutes as a mixture of a major monomeric species and minor dimeric species, whereas TmDFLP elutes mostly as a dimer, with addition of higher-order oligomers (Figure 3). Higher propensity of TmDFLP to form dimers might be a consequence of a disulfide bridge formation between the protomers, absent in CaDFLP, as discussed below.

**Figure 3.**
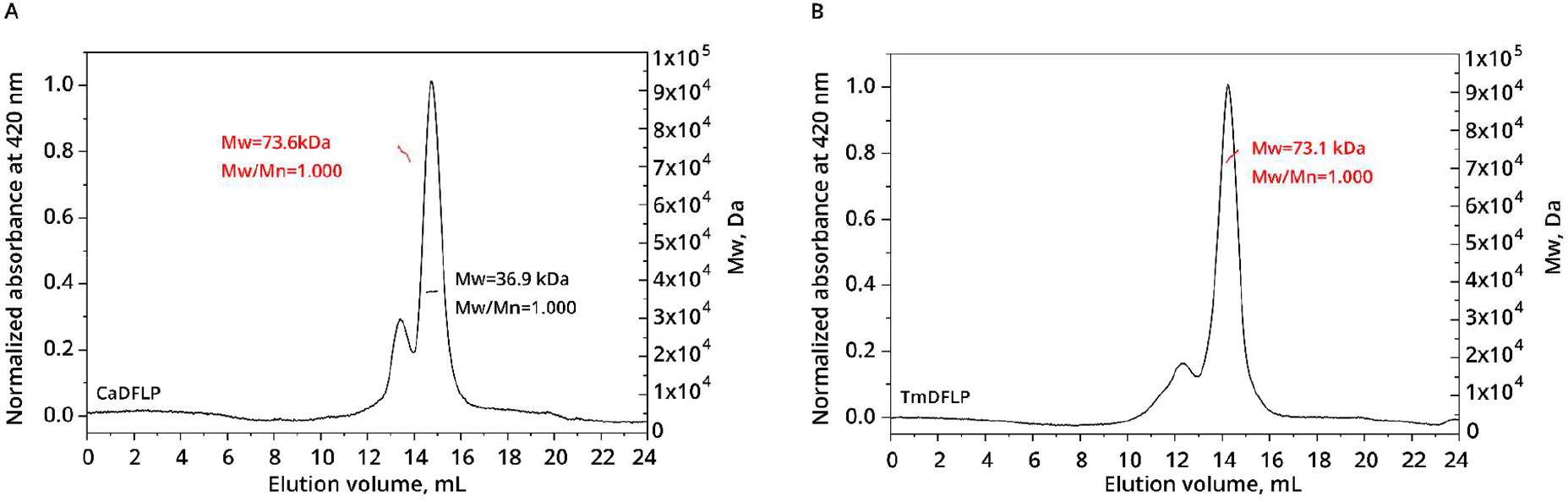
SEC-MALS analysis of DFLP proteins. A) Chromatogram for CaDFLP. B) Chromatogram for TmDFLP. The theoretical molecular weight (Mw) of DFLP monomers and dimers (with added purification tags) is approximately 36 and 72 kDa, respectively.

### Crystallographic structure of TmDFLP

To gain further insight into the structural properties of DFLPs, we sought to determine their crystal structures. Crystallization of CaDFLP was unsuccessful, whereas TmDFLP formed crystals that diffracted anisotropically with resolution cutoffs along the ellipsoid axes of 2.39, 2.84 and 3.13 Å (Table S1). Crystal structure revealed four polypeptide chains in the asymmetric unit, comprising residues 2 to 289, 2 to 287, 2 to 288 and 2 to 291 for chains A, B, C and D, respectively, organized as two symmetrical dimers. Interdomain linker residues 140-151, 141-151, 141-152 and 141-152 for chains A, B, C and D, respectively, were not resolved, presumably due to flexibility. The four TmDFLP molecules are highly similar with all atom RMSD between the chains ranging from 0.54 to 1.06 Å. All atom RMSD between the two dimers is 0.94 Å and backbone atom RMSD is 0.69 Å. All atom RMSD values between the individual protomers and the monomeric AlphaFold2 model available from UniProt are between 0.80 and 1.21 Å.

The structure of a TmDFLP dimer is presented in Figure 4. The two protomers are linked covalently via a disulfide bridge formed by cysteines 184, the only cysteines present in TmDFLP (Figure 4A). Similarly located cysteines are present in 19% of the proteins within the DFLP family and are absent in CaDFLP (see also the sequence alignment below). NTD and CTD of each protomer form an antiparallel arrangement similar to that found in bacterioferritins (Figure 4C,D). NTD and CTD, which have sequence identity of only 23.5%, align well, with backbone RMSD of α-helical regions of 0.79 Å.

**Figure 4.**
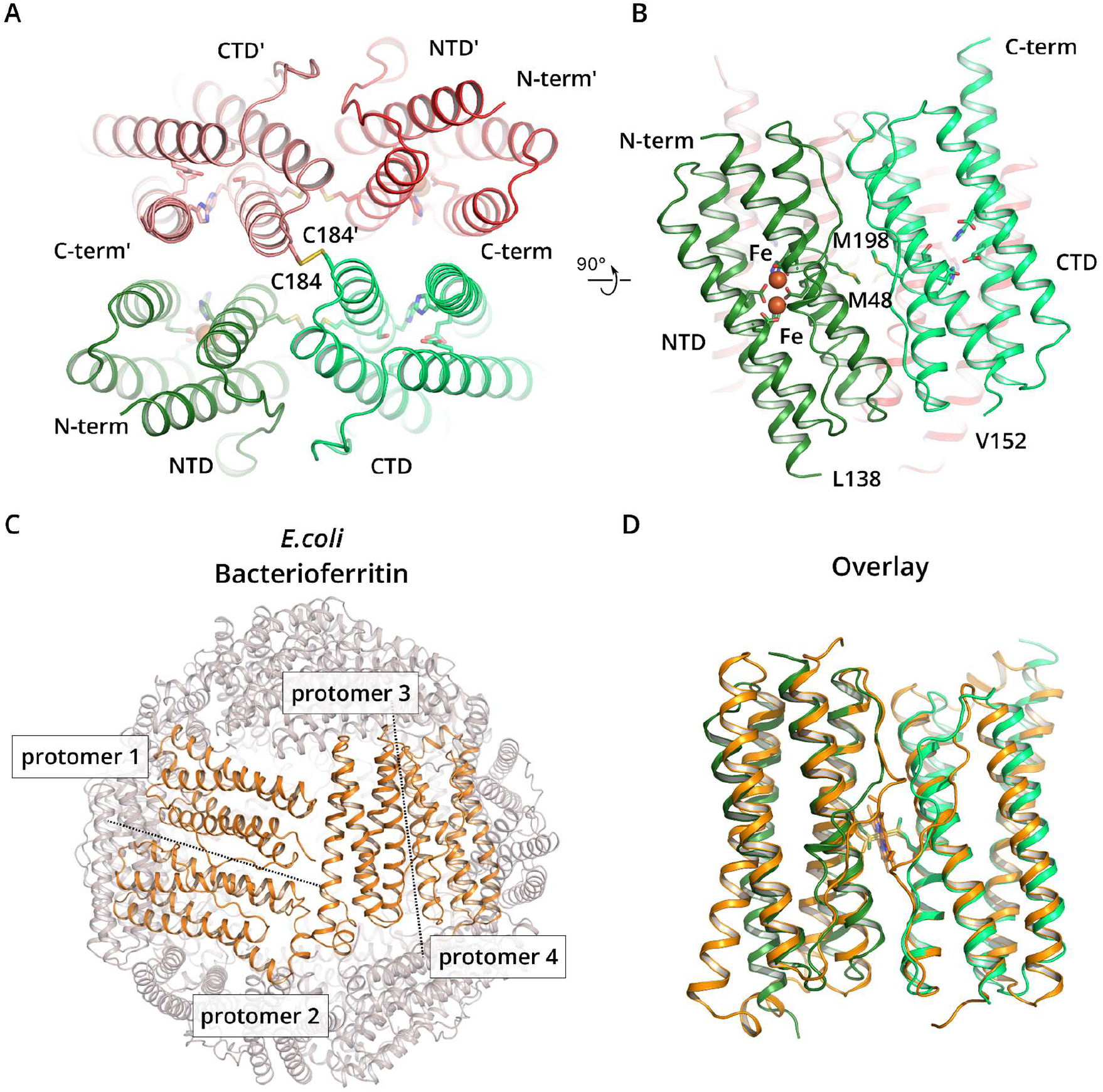
Crystal structure of TmDFLP. The protein is a parallel symmetric dimer. The relative arrangement of NTD and CTD is similar to that of the protomers in classic ferritins, but arrangement of the domains from different protomers is novel. A) View along the symmetry axis of the dimer with the protomers shown in shades of green and red. Cysteines 184 form a disulfide bridge. B) Side view of the dimer. M48 and M198 form a putative heme-binding site, although no heme is observed. Two putative iron-binding sites (orange) are observed in NTD. The loop connecting NTD and CTD (residues L138-V152) is not resolved in the electron density maps. C) Relative orientation of protomers in the bacterioferritin nanocage (PDB ID 1BFR^[25]^). Four adjacent protomers within the 24-mer are shown in orange for convenience. D) Structural superimposition of TmDFLP protomer (green) with *E. coli* bacterioferritin (orange, PDB ID 1BFR^[25]^). Heme and coordinating methionines (in the middle) are shown using sticks.

While the color and absorption spectrum of purified TmDFLP in solution signaled presence of heme, no heme was observed in the crystal structure. At the same time, heme can be readily placed in between NTD and CTD when the protein is modeled in AlphaFold3 (Figure S1); the putative heme binding site is lined with hydrophobic amino acids, there are no obvious steric clashes, and both methionine residues that could ligate the heme’s iron as observed in bacterioferritins (M48 and M198) are present (Figures 4B and S1). Moreover, these methionines are 100% conserved among DFLPs (see below). Thus, we assume that DFLPs may bind heme similarly to classic bacterioferritins, but heterologously produced proteins bound sub-equimolar amounts of heme due to suboptimal expression or purification conditions, and only the apo form of TmDFLP formed crystals.

### SAXS study of TmDFLP in solution

To see whether the higher-order oligomers or aggregates sometimes detected in TmDFLP preparations might correspond to partially formed ferritin-like spherical assemblies, we performed a small angle X-ray scattering experiment for different SEC fractions (SEC-SAXS) of a metal affinity-purified sample (Table S2). SEC profile demonstrated a peak corresponding to dimers at ∼14 ml, a peak corresponding to aggregates at 8 ml (void volume), and a set of unresolved intermediate peaks (Figure 5A). We selected six ranges (I to VI) for further detailed consideration (Figure 5A, Table S2d), with the range VI corresponding to dimeric TmDFLP. Plateauing of the radius of gyration (Rg---) in the range VI indicates homogeneity of the eluted sample in this range. CRYSOL fit^[26]^ obtained using the crystal structure of the TmDFLP dimer is in a satisfactory agreement with the experimental SAXS data merged over the range VI (Figure 5B and Table S2e).

**Figure 5.**
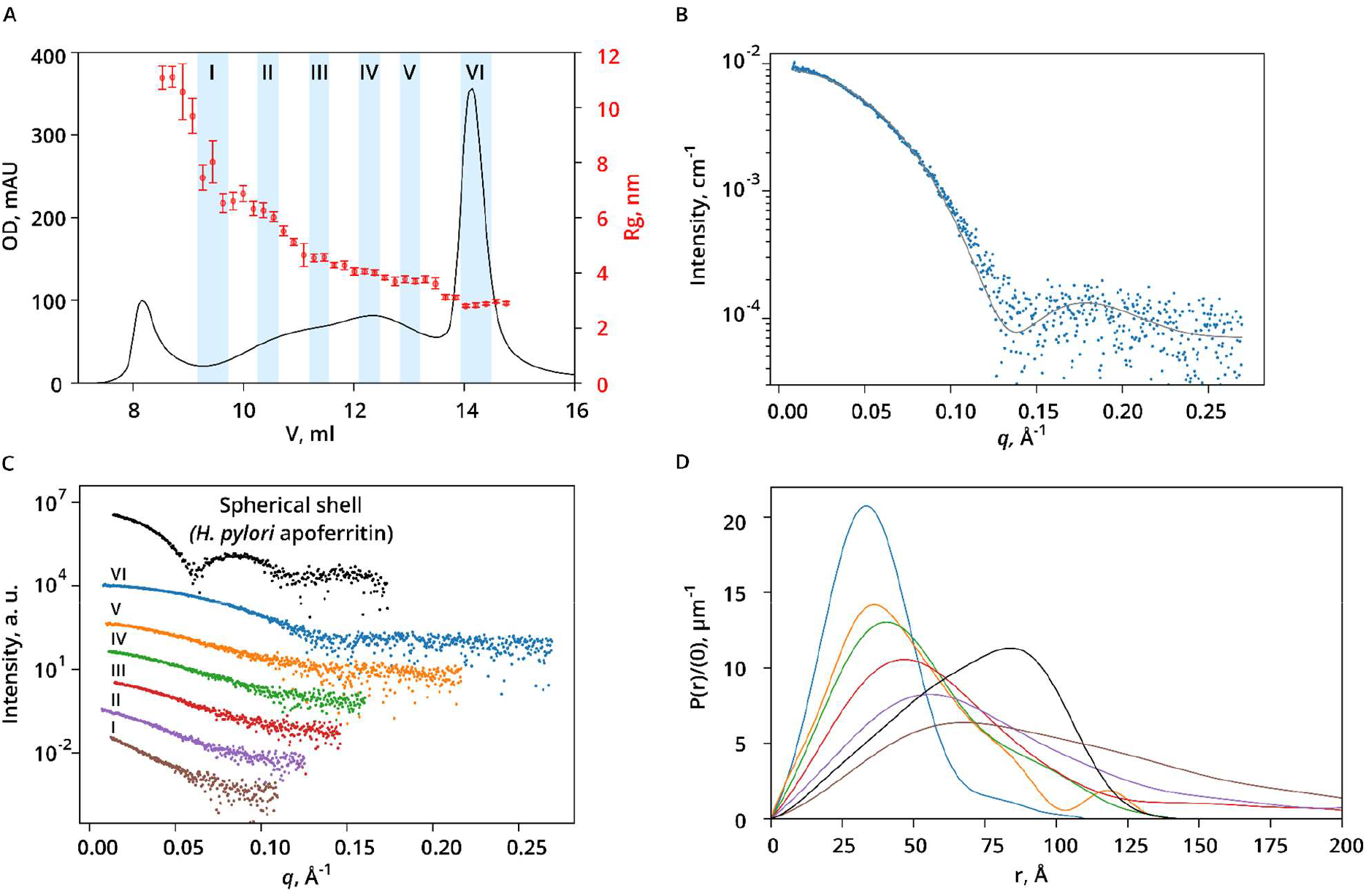
SEC-SAXS analysis of TmDFLP oligomers. A) SEC-SAXS profile of TmDFLP obtained using 24 mL Superdex 200 GL column. Black curve corresponds to UV absorption at 280 nm (left ordinate axis) and red signs correspond to the Rg values calculated from Guinier approximation (mean values and uncertainties arising from data fitting, right ordinate axis). Ranges selected for detailed analysis (I-VI) are highlighted in blue. B) SAXS curve for the range VI (blue dots) and CRYSOL fit obtained using the crystal structure of TmDFLP dimer (black line). C) SAXS curves merged over the selected ranges. SAXS curve for a shell-forming *H. pylori* apoferritin (SASBDB ID SASDTW4) is provided as a reference (black). D) Pair distance distribution functions normalized to intensity at zero angle I(0), derived from the SAXS curves presented in panel C. None of the curves corresponding to the TmDFLP sample displays a characteristic peak at ∼85 Å.

To check whether any of the remaining oligomers in the ranges I-V correspond to particles with a near-spherical shell form, we compared the respective experimental SAXS curves and pair distance distribution functions P(r) with the data for *H. pylori* apoferritin^[27]^ (SASBDB ID SASDTW4) (Figure 5C,D). For each range, the shapes of the scattering curve I(*q*) and of the pair distance distribution function P(r) qualitatively differ from those of the ferritin shell, indicating differences in the respective molecular structures (Figures 5D and S2). In particular, position of the maximum in P(r) is below half of the maximum pair distance for the TmDFLP fractions, which is characteristic for elongated particles and not for hollow or solid spheres. We also found that SAXS data for ranges III-V can be well approximated by mixtures of dimeric, tetrameric and octameric TmDFLP assemblies built based on crystal contacts (Figure S3 and Table S2f). Based on these data, we conclude that TmDFLP does not assemble into a spherical shell globule, while complexes with higher stoichiometry may correspond to oligomers with interfaces observed in crystals.

### AlphaFold modeling of DFLPs and related proteins

As the next step, to see whether DFLPs could theoretically form higher-order shell-like assemblies, we attempted to model various oligomers of TmDFLP and CaDFLP using AlphaFold3^[28]^. Because no DFLP structure was available in the Protein Data Bank at the moment of AlphaFold3 training, the algorithm was not expected to display any bias or memorization that could produce dimers rather than nanocage-like oligomers. AlphaFold3 confidently predicted correctly assembled nanocages for well-known ferritins and bacterioferritins, yet no sphere-like structures could be obtained for DFLPs. On the other hand, AlphaFold3 confidently predicted symmetric dimer models for DFLPs.

We also modeled representative proteins from the clusters that are situated in between DFLPs and bacterioferritins on the phylogenetic tree (Figure 1) and observed either dimers or dodecameric assemblies similar to PDB ID 2QQY (Figure 6). These dodecamers are similar to recently described mini-bacterioferritins^[29]^, which form a clade adjacent to bacterioferritins (Figure 1). Interestingly, we also identified a small cluster of DFLP-related proteins with a single ferritin-like domain. These single ferritin-like domain proteins are confidently modeled as heme-bound DFLP-like dimers of dimers by AlphaFold3 and thus might have evolved from DFLPs.

**Figure 6.**
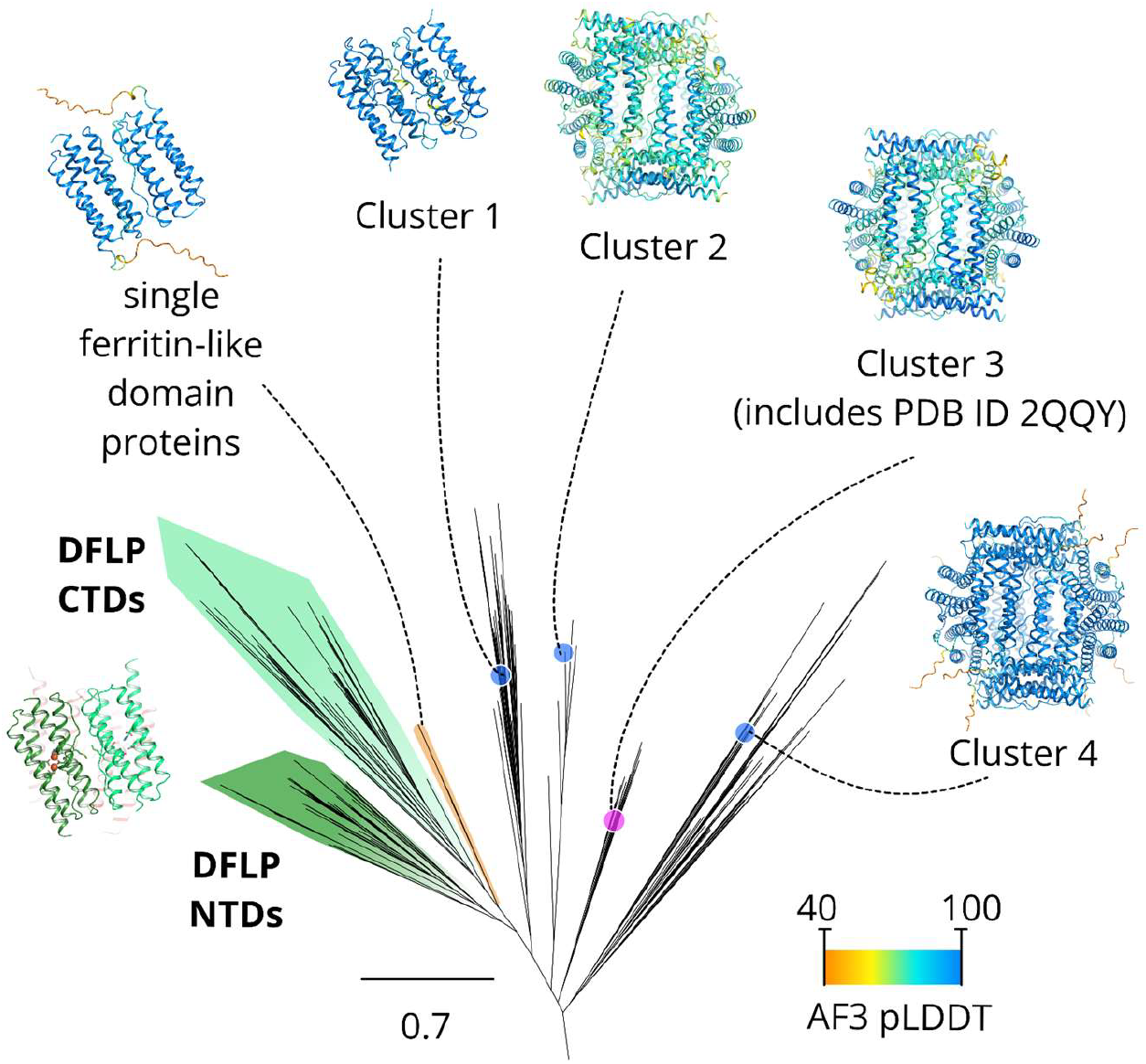
AlphaFold models of representative proteins from the bacterioferritin and DFLP-related clusters. Representatives from cluster 1 can only be modeled reliably as symmetric dimers and not as nanocages. Representatives from clusters 2-4 are modeled as 12-meric nanocages that are similar to PDB ID 2QQY^[22]^ (a structurally characterized member of cluster 3 of unknown function) and mini-bacterioferritins^[29]^.

### Ferroxidase sites of DFLPs

Given that TmDFLP is related to ferritins, it is informative to inspect putative diiron ferroxidase active sites in its two domains (Figure 7). NTD displays a classic bacterioferritin ferroxidase active site with four glutamates (E17, E47, E92 and E122) and two histidines (H50 and H125) coordinating two ligands (Figure 7A). Electron densities do not support placement of iron atoms with full occupancies; we modeled the ion close to E17 at the occupancy of 75% and the other one at the occupancy of 25%. Incomplete occupancy of a diiron site is a common occurrence in the literature^[30,31]^ and we presume that this site is functional. The iron atoms could be partially lost during purification, due to intrinsic protein ferroxidase activity, or during crystallization, especially due to the low pH value of the precipitant solution of 4.2. CTD of TmDFLP displays an uncharacteristic motif with three glutamates (E167, E240 and E270) and one serine (S197), whereas both histidines characteristic for bacterioferritins are present (H200 and H273) (Figure 7B). No clear densities that could be ascribed to any ligand are observed there. Consequently, at present, we cannot unequivocally conclude whether this site may display any ferroxidase activity. Analysis of the all identified DFLPs reveals that the motif comprising four glutamates and two histidines (EEHEEH, in the order of appearance in the sequence) is conserved in 79.6% of NTDs and only in 57.3% of CTDs, with the most common deviations being ESHEEH (13.3%) and KKQVEQ (8.9%). Overall patterns of amino acid conservation differ between the two DFLP domains (Figure 7C,D), with CTD being more variable, as also indicated by stronger divergence in the CTD clade in the phylogenetic tree (Figure 1).

**Figure 7.**
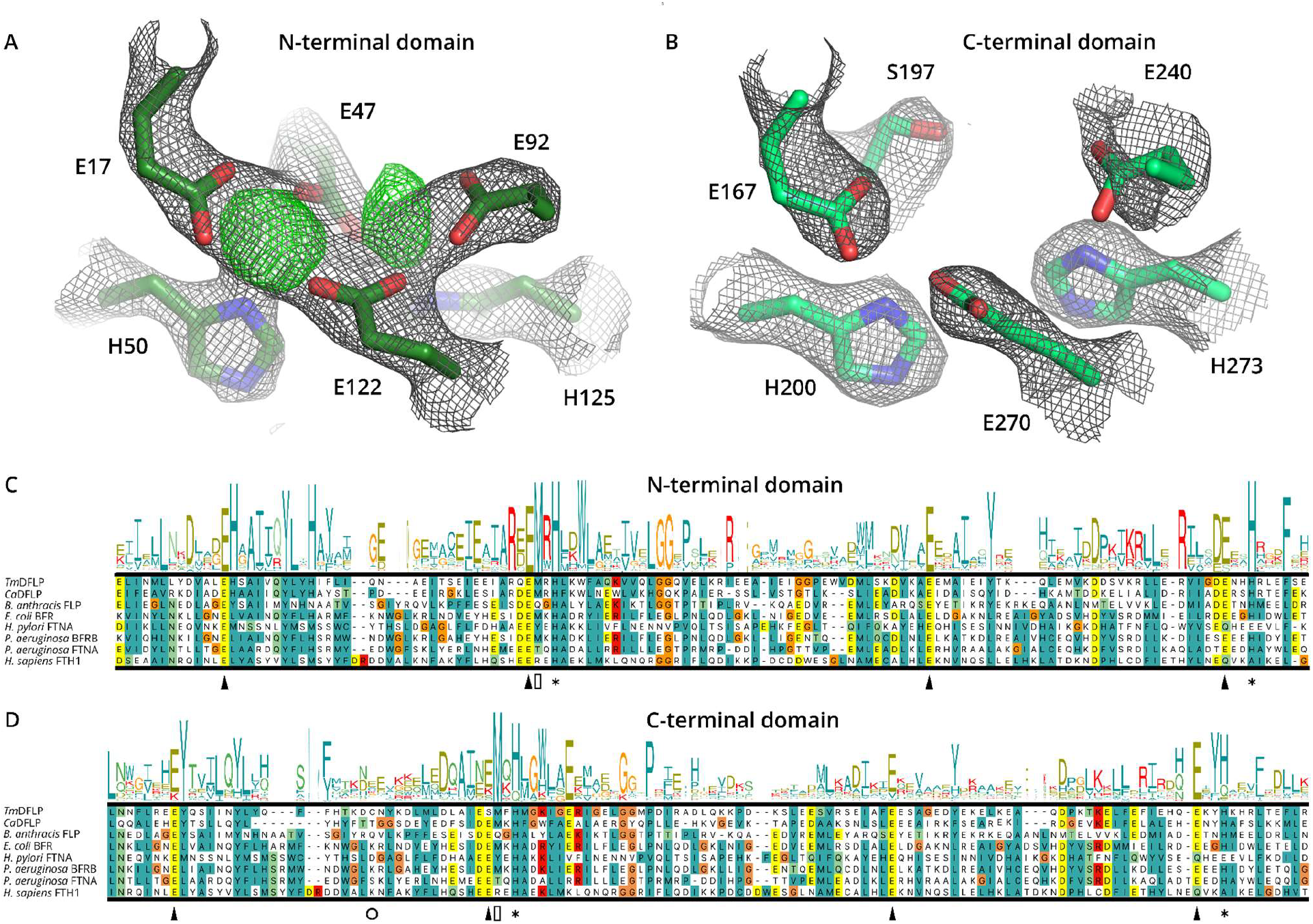
Putative ferroxidase sites in DFLPs. A) and B) 2Fo-Fc Electron density maps (black) for tentative active sites in NTD (A) and CTD (B) of TmDFLP. The maps are overall comparable in the four protomers in the crystallographic asymmetric unit, and only the chain A models and densities are shown. In NTD, difference densities in the two putative iron positions support placement of iron atoms with occupancies of 0.75 and 0.25, respectively, or an iron atom with an occupancy of 0.75 and a water molecule with an occupancy of 1.00. In CTD, no clear densities that could be ascribed to any ligand are observed. 2Fo-Fc electron density maps are contoured at the level of 1.8 σ in A and 1.2 σ in B for convenience. Difference electron Fo-Fc density map (green) prior to placement of ligands is contoured at the level of 3σ. C) and D) Representative sequences of ferritin family members compared with Weblogo plots and sequences for NTD (C) and CTD (D) of DFLPs. Conserved glutamate, histidine and methionine positions are marked with triangles, asterisks and rectangles, respectively. Position of the cysteine forming an intra-protein disulfide bridge at the dimerization interface in TmDFLP is marked with a circle. CTDs and their possible ferroxidase site residues are less well conserved compared to those of NTDs.

### Ferroxidase activity of DFLPs

Given that ferritins and ferritin-like proteins may oxidize iron, we proceeded with characterization of the possible catalytic activity of DFLPs. Due to lower activity of CaDFLP observed in preliminary tests and because of its lower expression level, we focused on TmDFLP. First, we assessed the ability of TmDFLPs to remove Fe^2+^ from solution using a free Fe^2+^ assay. We added FeSO_4_ to DFLP samples stepwise and examined the time dependence of the free Fe^2+^ concentration in the solution. The free iron content rapidly decreased in presence of TmDFLP, while remaining almost constant in the control sample (Figure 8A). Interestingly, we observed a decline of TmDFLP activity with time: each new Fe^2+^ portion was processed slower. The protein is negatively charged at neutral pH (62 Asp and Glu amino acids versus 40 Arg and Lys), and the drop in its activity might be the result of inhibition of TmDFLP by Fe^3+^ ions that repel Fe^2+^. Measurements of Fe^2+^ and Fe^3+^ concentrations following addition of Fe^2+^ to the sample clearly demonstrate the iron oxidation activity of TmDFLP (Figure 8B). Combined with structural data (Figure 7), these findings unequivocally characterize the protein as a ferroxidase.

**Figure 8.**
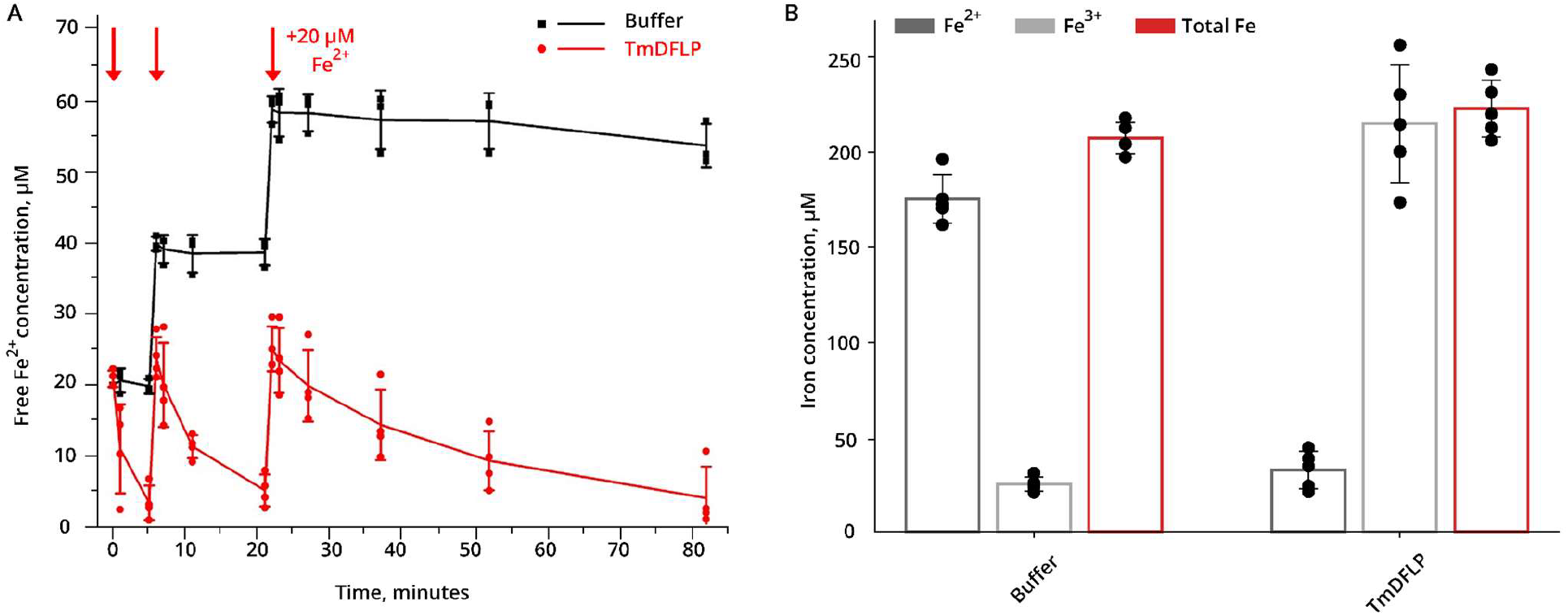
Ferroxidase activity of TmDFLP. A) Time dependence of free Fe^2+^ concentration in the buffer solution (control sample) and in presence of 10 μM of TmDFLP protomers. Addition of 20 μM iron doses is indicated with red arrows. The sample was incubated at 20 °C. B) Fe^2+^, Fe^3+^ and total iron concentrations in the control sample and in the solution containing 50 μM of TmDFLP protomers, following addition of 200 μM FeSO4 and 45 min incubation at 25 °C.

### Interaction of DFLPs with encapsulin

To better understand the function of DFLPs in the native hosts, we proceeded towards analyzing their genomic environment. For example, the genes of previously studied bacterioferritins are neighboring the genes of bacterioferritin-associated ferredoxin, which is necessary for facilitation of the ferric ion release^[6]^. We could not identify any genes encoding iron metabolism proteins at the positions from +6 to −6 relative to the TmDFLP gene. In contrast, the neighborhood of the CaDFLP gene includes several iron binding-associated genes. A gene encoding a protein with rubredoxin domain is located in the −2 position (WP_073257887.1), and a similarly oriented gene for a protein with rubrerythrin domain is observed in the −3 position (WP_073257889.1). Such arrangement may point to origin from a single gene containing both domains, as observed in classic rubrerythrins, which are commonly found in other organisms^[32]^. Additional analysis revealed another rubrerythrin gene in the +4 position (WP_073257876.1), and a ferredoxin-coding gene in the −6 position (WP_073257895.1). We note that distance in 6 open reading frames is not sufficient to make a confident claim about functional relations between ferredoxin and CaDFLP.

On the other hand, we identified an encapsulin-coding gene, WP_084098969.1, in the position −1 relative to the CaDFLP gene. The main function of encapsulins is packaging of proteins in nanocompartments^[33–35]^. Importantly, some encapsulins were previously shown to harbor ferritin-like proteins, thus playing a role in iron metabolism^[17,19,20]^. Moreover, previous studies noted presence of DFLPs in encapsulin operons, but assumed that they were nanocage-forming bacterioferritins and did not follow up with experimental verification^[33,36,16]^.

*C. acetigignens* protein encoded by WP_084098969.1 is annotated as Type 1 encapsulin shell protein in InterPro (IPR007544) and as Encapsulating protein for peroxidase in Pfam (PF04454). Currently, encapsulins are divided into four families depending on sequence similarity, encapsulin binding motif position in cargo proteins and the operon structure^[37]^. Among them, the most extensively studied groups with experimentally resolved encapsulin structures are Family 1 and Family 2. One of the main differentiating factors between the Family 1 and Family 2 is the order of the encapsulin and cargo genes in operon: Family 1 encapsulin genes are located downstream of the cargo, while in Family 2 they are located upstream. The second key difference is location and sequence of the encapsulin-binding fragment in cargo proteins. Typically, the binding motif is placed at the end of the cargo that is closer to the encapsulin gene: Family 1 cargoes have the sequence at the C-terminus of the protein, and Family 2 cargoes at the N-terminus^[36]^.

Taking into account the InterPro and Pfam annotation, and also the location of the *C. acetigignens* encapsulin gene downstream of CaDFLP gene, we assert that WP_084098969.1 belongs to the encapsulin Family 1, and that CaDFLP serves as a core cargo (Figure 9A). At the same time, *T. minervae* possesses an encapsulin gene in another part of the genome, thus TmDFLP might serve as a secondary cargo (Figure 9A). Analysis of the genomic environment of other DFLPs reveals 3 cases of reversed gene order, with the encapsulin gene located upstream of the cargo, which is uncharacteristic for Family 1 encapsulins (Figure 9A)^[36]^. Interestingly, the single ferritin-like domain clade (Figure 6) contains three members with encapsulin-targeting peptides and encapsulin genes downstream (Figure 9A).

**Figure 9.**
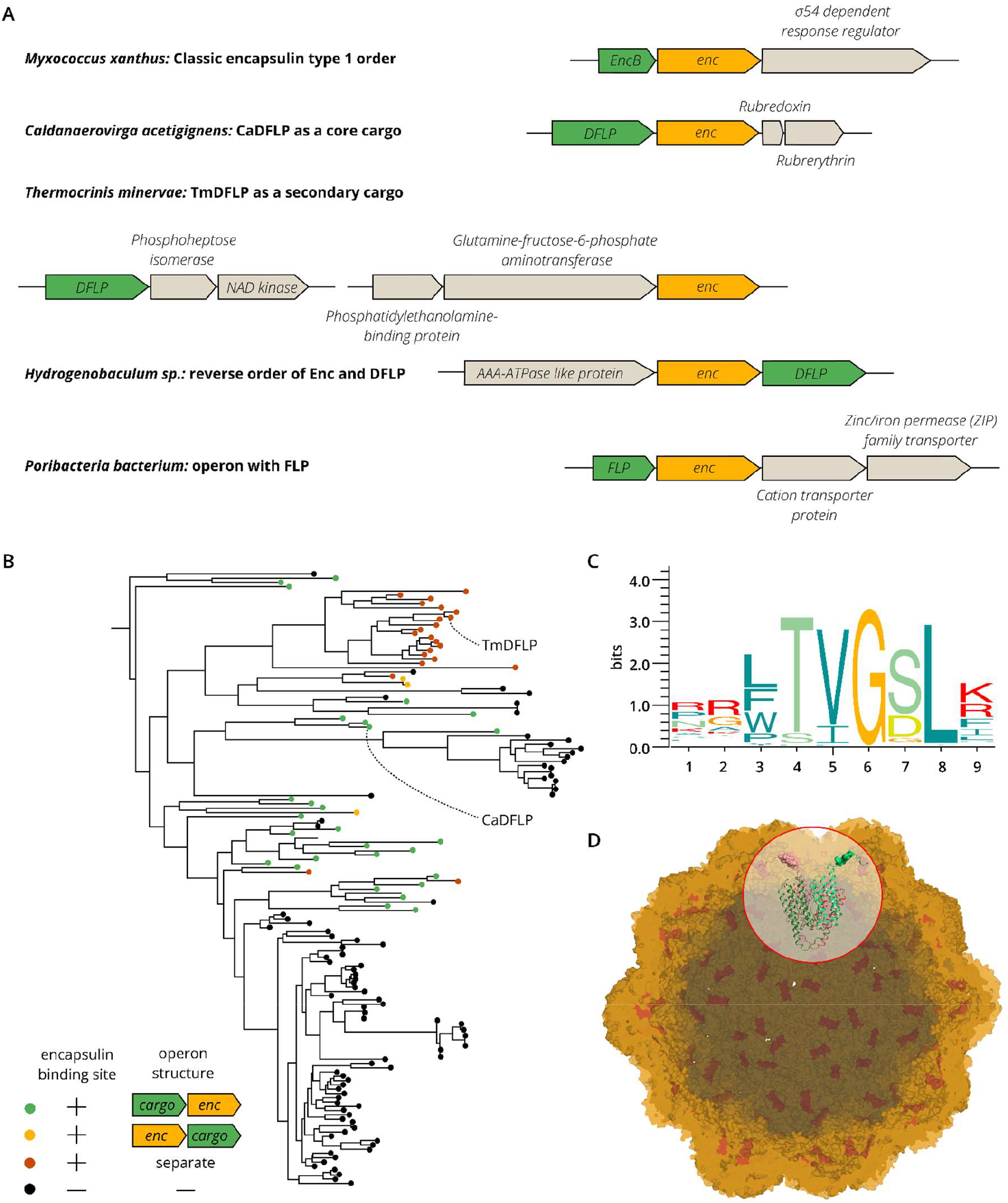
DFLP-encapsulin interaction analysis. A) Genomic environment for the investigated proteins. CaDFLP gene is located upstream of the encapsulin gene similarly to the typical encapsulin type I operon in *Myxococcus xanthus*. The TmDFLP gene and the encapsulin gene are separated from each other in the genome. Other genomic environment types in the investigated cluster include reversed order of encapsulin and DFLP genes, and also operons with FLP genes containing single ferritin domain. B) Phylogenetic tree of DFLPs with encapsulin interactions marked. C) Sequence logo for encapsulin-binding motifs in DFLPs (see also Figures S5 and S6). D) Model of a CaDFLP dimer within an encapsulin shell with a symmetry T=3 (based on *M. xanthus* EncA, PDB ID 8VJO, the most closely related structurally characterized encapsulin). DFLP protomers are shown in green and red with TPs visualized in surface mode. Remaining TPs are shown in red.

As the next step, we analyzed the possible DFLP-encapsulin interaction from the structural viewpoint. Several targeting peptide (TP)-encapsulin complexes have been structurally characterized previously. The respective sequence motifs are LGI(RK) in *Thermotoga maritima* (PDB ID 3DKT^[38]^), LGIGSL in *Haliangium ochraceum* and *Rhodospirillum rubrum* (PDB IDs 5N5F^[18]^ and 5DA5^[17]^), LTVGSL for EncB, EncC and EncD proteins in *Myxococcus xanthus* (PDB IDs 7S5K^[20]^, 7S8T^[20]^ and 8VJO^[39]^) and FTVGSL in *Quasibacillus thermotolerans* (PDB ID 6N63^[19]^). Thus, the general TP motif may be described as [LF]-x-[VI]-x-x-[LI]^[40]^, with the minimal required sequence of [LF]-x-[VI]-x, based on shortened TP in *Thermotoga maritima*^[41]^. We used the ScanProsite tool^[42]^ to search for [LF]-x-[VI]-x-x-[LI] anywhere in the sequence of CaDFLP and found a sequence FTVGSL at the C-terminus of the protein. Moreover, we found a similar TP site FTIGGL at the C-terminus of TmDFLP, although the TmDFLP gene does not belong to a Family 1 operon. Structural alignment of AlphaFold models of encapsulin-DFLP peptide complexes from *C. acetigignens* and *T. minervae* with the experimentally resolved structure of *M. xanthus* EncA-EncB TP complex (PDB ID 7S5K) reveals close correspondence between the binding interfaces (Figure S4).

We then performed a similar analysis of the remaining 155 proteins in the DFLPs cluster using an iterative procedure, where AlphaFold modeling was performed to identify the encapsulin-binding fragments, update the motif definition and search for the putative binding sites in the rest of the sequences. As a result, existence of probable encapsulin-binding motifs was confirmed for 56 DFLPs, including CaDFLP and TmDFLP. Based on the analysis, distinct DFLP groups can be identified in the DFLP clade of the phylogenetic tree (Figure 9B). Proteins grouping with TmDFLP have an encapsulin binding site but lack encapsulin genes in the closest genomic environment, and thus are proposed to play a secondary cargo role. Others, like CaDFLP, have conserved binding sites and have Type I encapsulin-like operon structure, thus probably playing a core cargo role. Yet for others, the order of the genes is reversed, while the TP motifs are still found at the C-termini of the proteins. Finally, some DFLPs lack both encapsulin binding site and encapsulin gene in the genomic environment; possible function of these proteins is not clear at present.

Performed analysis allows us to better describe the encapsulin binding motif for FLP proteins (Figure 9C). In accordance with previous studies^[17,18,20,35,39]^, positions 3, 5 and 8 are occupied with hydrophobic amino acids (isoleucine, leucine, or valine), which mediate interactions with encapsulin, and positions 4, 6 and 7 are occupied with amino acids that may provide flexibility (Figures 9C, S5 and S6). Highly conserved glycine residue was observed at the position 6 in all 56 studied DFLP proteins, as well as in the 6 structurally characterized complexes, which indicates its essential role. However, prevalence of glycine and serine residues in positions 1 and 2, assumed to be crucial for additional flexibility, is not detected for DFLPs. Finally, as noted in earlier analyses^[35]^, the encapsulin-binding motif in DFLPs may be followed by additional 1 to 14 C-terminal residues (Figure S6).

## Conclusion

Here, we identified and comprehensively characterized a group of bacterioferritin-related proteins containing duplicated four-helical ferritin domain, DFLPs. Our analysis shows that DFLPs likely descended from bacterioferritins, keeping some of their properties (heme binding and ferroxidase activity), but lacking the others (formation of nanocages and iron storage). At least some DFLPs are targeted into encapsulin nanocompartments, which could thus serve as iron-storing nanocages that are alternative to classic 24-mers formed by ferritins, a notion further supported by the lack of any other ferritin domain-containing proteins in the respective organisms. Overall, reported findings extend the available knowledge of FLPs with diiron sites and of encapsulin systems, and highlight the power of combining experimental and bioinformatic approaches.

## Methods

### Data collection and clustering

Protein sequences were downloaded from InterPro v. 98.0^[21]^ (released on January 25th, 2024). Domain architecture was determined by using AlphaFold2^[43]^ (ColabFold^[44]^ v. 1.5.5 with alphafold2_multimer_v3 weights), AlphaFold3^[28]^ and HMMER^[45]^. The analyzed set contained the following three groups: i) The group of sequences where a full or partial ferritin domain is detected (but not two or more), clustered using Usearch 11.0.667^[46]^ with identity percentage set at 35%. ii) The group of sequences where two ferritin domains are detected. These sequences were aligned using MUSCLE v. 5.15.2^[47]^ and divided into N-terminal domains and C-terminal domains using the domain border in the *T. minervae* protein (UniProt ID A0A1M6SI41) as a reference (residues 142-143). iii) The group of sequences with experimentally determined structures, as downloaded from InterPro.

### Phylogenetic tree

Sequence alignment was performed using HMMER with parameters [-hmmalign --trim]. The alignment was manually reviewed and nine entries were removed as they contained inserts in the positions where more than 99% sequences had gaps. The phylogenetic tree was built using Fasttree 2.0^[48]^, and annotation was performed in Figtree v.1.4.4^[49]^ (Rambaut A. 2018. FigTree v.1.4.4).

### Analysis of the genomic environment

GenBank IDs for genes and assemblies were obtained using the IDmapping tool in UniProt^[50]^.

### Cloning, protein expression and purification

The nucleotide sequences encoding CaDFLP (gene BUB66_RS09395, UniProt accession code A0A1M7LGY2) and TmDFLP (gene SAMN05444391_1032, UniProt accession code A0A1M6SI41) with C-terminal 6×His tags were codon-optimized for expression in *E. coli* and introduced into the pET-28a expression vector (Novagen, Merck, Germany). Each protein was expressed in *E. coli* strain BL21 Gold (DE3). Cells were cultured in shaking baffled flasks in ZYP-5052 autoinduction medium^[51]^ containing 50 μg/mL kanamycin at 30 °C. Harvested cells were resuspended in a buffer containing 50 mM NaCl, 20 mM Tris, pH 8.0, with addition of 0.5 mM PMSF and disrupted by M-110P Lab Homogenizer (Microfluidics, USA). The lysate was clarified by removal of cell membrane fraction by ultracentrifugation at 100,000 g for 1 h at 10 °C. The supernatant was incubated with Ni-nitrilotriacetic acid (Ni-NTA) resin (Qiagen, Germany) on a rocker for 1 hour at 4 °C. The Ni-NTA resin and supernatant were loaded on a gravity flow column. The protein was eluted by a buffer supplemented by 200 mM imidazole. The eluate was concentrated and subjected to a size-exclusion chromatography (SEC) on Superdex® 200 Increase 10/300 GL column (GE Healthcare Life Sciences, USA) in a buffer containing 50 mM NaCl, 20 mM Tris, pH 8.0. Protein-containing fractions were pooled and concentrated for the following studies.

### Evaluation of iron removal kinetics

We evaluated the changes in free Fe^2+^ concentrations similarly to Ref. ^[52]^. Briefly, 10 mM FeSO_4_ stock solution was added to the samples in three steps, each time 1/500 v/v to increase the concentration of Fe^2+^ by 20 μM FeSO_4_, followed by thorough mixing and incubation at 20 °?. For measurements, 100 μL of the sample solution was collected and mixed with ferrozine (final concentration of 0.24 mg/mL) in a 96 well plate. The amount of free Fe^2+^ in collected probes was calculated from the optical density at 560 nm, measured using Synergy H4 Hybrid Microplate Reader (BioTek, USA). Calculation of iron concentration in the probes was performed using a calibration curve obtained for probes with 10-100 μM FeSO_4_. The control sample contained 50 mM NaCl, 20 mM Tris, pH 6.8, and the test sample additionally contained 10 μM of the purified protein. In order to avoid iron autooxidation, 10 mM FeSO_4_ stock solution was prepared immediately before running each assay.

### Evaluation of iron oxidation

To study iron oxidation during long term incubation, we added 10 mM FeSO_4_ stock solution to the samples to reach the final concentration of 200 μM. Iron was added in two steps of 100 μM, with a pause of 10 minutes between the doses. The incubation, conducted at 25 °C, was stopped by addition of a three-fold volume of acidic buffer (50 mM NaCl, 20 mM Tris, 0.5% HCl) 45 minutes after the start of the experiment. After that, the protocols varied for each measurement.

To determine the concentration of Fe^2+^, the samples were incubated for 5 minutes at 90 °C. Ammonium acetate was added to reach the final concentration of 0.25 M, and 100 μL of solution was mixed with ferrozine for the spectrophotometric assay described above.

To determine the concentration of Fe^3+^, the samples were incubated for 10 minutes at 90 °C. Then, the sample was mixed with potassium thiocyanate to reach a final concentration of 1%, in accordance with Ref. ^[53]^. The amount of Fe^3+^ in collected probes was calculated similarly to the protocol used for Fe^2+^, based on the optical density at 480 nm. Calculation was performed using a calibration curve obtained for probes with 10-100 μM of Fe_2_(SO_4_)_3_.

To determine the concentration of total iron (both Fe^2+^ and Fe^3+^), in accordance with Ref. ^[54]^, potassium permanganate was added to the samples right before the incubation at 90 °C to reach a final concentration of 0.75%. After 15 min of incubation, ascorbic acid and ammonium acetate were added to reach the final concentrations of 0.1 M and 0.25 M, respectively, and 100 μL of solution was mixed with ferrozine for the spectrophotometric assay described above.

### Crystallization of TmDFLP

The protein sample was concentrated to 4 mg/mL prior to crystallization. Sitting drop vapor diffusion crystallization trials were set up using an NT8 robotic system (Formulatrix, USA) and stored at 20 °C. The drops contained 200 nL protein solution and 200 nL reservoir solution. The best crystals were obtained using the following solution as the precipitant: 40 % v/v Ethanol, 5 % w/v PEG 1000, 0.1 M Phosphate/citrate buffer, pH 4.2. Stick-shaped crystals appeared and reached the final size of ∼100-300 μm within a month. The crystals were harvested using micromounts, then flash-cooled and stored in liquid nitrogen.

### Diffraction data collection and processing

The diffraction data were collected at 100 K at Shanghai Synchrotron Radiation Facility (SSRF) BL17UM High Performance Membrane Protein Crystallography Beamline. Diffraction images were processed using XDS^[55]^. AIMLESS^[56]^ was used to merge and scale the data. The data collection and processing statistics are presented in Table S1.

### Structure determination

The structure was solved using molecular replacement with Phaser^[57]^ and AlphaFold^[43]^ model of TmDFLP as a search model. The resulting model was refined manually using Coot^[58]^ and REFMAC5^[59]^.

### Multi-angle light scattering (MALS)

Absolute molecular mass of ferritins was measured by multi-angle static light scattering coupled with size-exclusion chromatography (SEC-MALS), which is independent of a protein shape and interactions with the chromatographic resin. CaDFLP and TmDFLP samples (100 μL each, ∼1 mg/mL), clarified by centrifugation for 5 min at 21,400g at 4 °C, were loaded using a Vialsampler (G7129A, Agilent) on a Superdex 200 Increase 10/300 column (Cytiva) pre-equilibrated by SEC buffer (20 mM Tris–HCl, pH 8.0, 150 mM NaCl) and run at a 1.0 mL/min flow rate using an Agilent 1260 Infinity II chromatography system equipped with a consecutive detectors – 1260 Infinity II WR diode-array detector (G7115A, Agilent), a miniDAWN detector (Wyatt Technology), and a refractometric detector (G7162A, Agilent, operated at 35 °C). The elution profiles were followed by absorbance at 420 nm specific to ferritin holoforms, by static light scattering intensity at two angles (90° and 135°) and by changes of the refractive index. The data were processed in Astra 8.0 software (Wyatt Technology) using the refractometric detector as a concentration source (dn/dc was taken equal to 0.186 for each protein). Normalization of the light scattering intensities at different angles detected by miniDAWN was carried out using a pre-run profile using a BSA standard (Wyatt). The polydispersity index (Mw/Mn) was close to 1.000 for all samples characterizing monodisperse particles.

### Small-angle X-ray scattering (SAXS)

Preliminary measurements were performed using SAXS instrument Rigaku MicroMax-007HF (MIPT, Dolgoprudny, Russia), previously described in Refs.^[27,60]^, and at the 4C SAXS II beamline at the Pohang Accelerator in Korea^[61]^. Scattering data presented in the manuscript were collected at beamline BL19U2 of National Facility for Protein Science Shanghai (NFPS) at Shanghai Synchrotron Radiation Facility (SSRF, Shanghai, China), described in Refs.^[62,63]^.

The wavelength (λ) of X-ray radiation was set as 1.03 Å. The sample-to-detector distance was set as 2.67 m (corresponding range of momentum transfer *q* = 4π sin(θ)/λ was 0.006-0.456 Å^-1^, where 2θ is the scattering angle). Scattered X-ray intensities were collected using a Pilatus3 2M detector (DECTRIS Ltd., Switzerland) with the exposure time of 1 second per frame.

SEC-SAXS was performed using the Superdex 200 GL column (24 ml, GE Healthcare, USA) with the flow rate set at 0.5 ml/min. The inline SEC was performed using commercial high-performance liquid chromatography (HPLC, Angilent 1260 Infinity, Angilent, USA). The buffer used was DPBS. The columns and the SAXS flow cell were maintained at 25 °C. The sample concentration was 9 mg/ml, and 100 μl sample was injected into the column after centrifugation.

The BioXTAS RAW^[64]^ and ATSAS^[65]^ software suites were used for data treatment, including PRIMUS^[66]^ for Guinier approximations, GNOM for calculation of the pair-distance distribution functions P(r), CRYSOL^[26]^ and OLIGOMER for fitting experimental data to atomic models and mixtures of known components, respectively. We also used methods of molecular weight determination, described in Refs. ^[67,68]^. See detailed information about the setup and analysis in Table S2.

## Data availability

Atomic coordinates and structure factors for the reported crystal structure have been deposited into the Protein Data Bank under the accession code 9VIW.

## Supporting data

Supporting Tables 1-2, Supporting Figures 1-6, and a Supporting Data file with a list of annotated DFLPs are available.

## Contributor Roles

Conceptualization: I.G. Funding acquisition: I.G., V.B. Investigation: A.A., A.R., I.N., A.Y., R.Al E., P.S., V.S., V.M., O.S., E.K., A.N., I.B., D.K., E.D., N.L., Y.R., N.N.S., Y.Y., V.B., A.V., S.V.B., I.V.M., I.G. Project administration: I.G., A.A., A.R. Writing – original draft: A.A., A.R., V.S., Y.R., N.N.S., I.G. Writing – review & editing: A.A., A.R., I.N., A.Y., R.Al E., P.S., V.S., V.M., O.S., E.K., A.N., I.B., D.K., E.D., N.L., Y.R., N.N.S., Y.Y., V.B., A.V., S.V.B., I.V.M., I.G.

## Acknowledgements

We acknowledge the Shanghai Synchrotron Radiation Facility in China for granting access to the synchrotron BL17UM beamline and thank the beamline staff for their exceptional technical assistance during our experiments. We thank the staff members of BL19U2 beamline (https://cstr.cn/31129.02.NFPS.BL19U2) at the National Facility for Protein Science in Shanghai (https://cstr.cn/31129.02.NFPS), for providing technical support and assistance in data collection and analysis. We thank Dr. Alexander I. Kuklin and Yury S. Semenov for their invaluable support with using the SAXS equipment Rigaku MicroMax-007HF at the Moscow Institute of Physics and Technology (MIPT, Russia). We also acknowledge the Pohang Accelerator Laboratory (PAL, Korea) for granting access to the 4C SAXS II beamline and thank the beamline staff for their hospitality and support.

Bioinformatic analyses and characterization were supported by the Ministry of Science and Higher Education of the Russian Federation (agreement 075-03-2025-662, project FSMG-2025-0003). Crystallographic data collection was supported by the Ministry of Science and Higher Education of the Russian Federation (agreement 075-03-2025-662, project FSMG-2025-0012). N.N.S. acknowledges that his work was partly supported by the program of the Ministry of Science and Higher Education of the Russian Federation.

