## Supporting Information for "Structural and Functional Characterization of Encapsulin-Targeted Double Ferritin Fold Ferroxidases"

Supporting Tables S1-S2

Supporting Figures S1-S6

**Supporting Table S1.** Crystallographic data collection and refinement statistics.

| | |
|---|---|
| **(a) Data collection** | |
| Space group | P 1 21 1 |
| **(b) Cell dimensions** | |
| a, b, c (Å) | 63.96, 105.16, 89.14 |
| α, β, γ (°) | 90.00, 90.27, 90.00 |
| Resolution (Å) | 46.50 – 2.45 (2.72 – 2.45) |
| <I/σI> | 5.1 (1.4) |
| CC1/2 (%) | 98.2 (59.8) |
| Completeness (%) | 65.1 (12.5) |
| Multiplicity | 6.9 (7.2) |
| Unique reflections | 28140 (1407) |
| **(c) Refinement** | |
| Resolution (Å) | 46.55 – 2.47 |
| No. of reflections | 27917 |
| $R_{work}/R_{free}$ (%) | 24.3/29.0 |
| **(d) No. of atoms** | |
| Protein | 9223 |
| Fe | 4 |
| Water and others | 5 |
| **(e) Average B factors (Å$^2$)** | |
| Protein | 40.4 |
| Fe | 43.6 |
| Water and others | 21.5 |
| **(f) R.m.s. deviations** | |
| Protein bond lengths (Å) | 0.0010 |
| Protein bond angles (°) | 0.5461 |
| **(g) Ramachandran analysis** | |
| Favored (%) | 98 |
| Allowed (%) | 2 |
| Outliers (%) | 0 |

**Supporting Table S2.** Experimental details of SAXS measurements and data evaluation summary.

| **(a) Sample details** | |
|---|---|
| | Ferritin-like protein from *Thermocrinis minervae* (TmDFLP) |
| Description of sequence | UniProt ID A0A1M6SI41 with N-terminal 6×His tag |
| Chemical formula | $C_{1595}H_{2505}N_{433}O_{488}S_{12}$ |
| Expected monomer mass (kDa) | 35939.76 |
| Approximate volume (Å$^3$) | 43484 |
| Partial specific volume v (cm$^3$ g$^{-1}$) | 0.7286 |
| Mean solute and solvent SLD ($10^{-6}$ Å$^{-2}$) | 9.40; 12.492 |
| Mean scattering contrast Δρ ($10^{-6}$ Å$^{-2}$) | 3.092 |
| Extinction coefficient $\varepsilon_{280nm}$ (M$^{-1}$ cm$^{-1}$) | 25900 |
| Sample concentration (mg ml$^{-1}$) | SEC-SAXS, injected 100 uL × 9 mg/ml |
| Solvent composition | Storage: 50 mM NaCl, 20 mM Tris, pH 8.0<br>SEC-SAXS: 137 mM NaCl, 2.7 mM KCl, 0.9 mM $CaCl_2$, 0.5 mM $MgCl_2$, 8.1 mM $Na_2HPO_4$, 1.5 mM $KH_2PO_4$, pH 7.4 |
| **(b) SAS data collection parameters** | |
| Instrument | BL19U2 Beamline with Pilatus3 2M (DECTRIS Ltd) (SSRF, Shanghai, China) |
| Wavelength (Å) | 1.033 |
| Beam geometry | 0.33 mm (H) × 0.05 mm (V);<br>sample-to-detector distance = 2.7 m |
| Sample configuration | Quartz capillary with 10 µm-thick walls and a 1.5 mm path length |
| q-measurement range (Å$^{-1}$) | 0.0064 – 0.4068 |
| q-scaling method | Calibration standard: silver behenate powder |
| Basis for normalization to constant counts | To transmitted intensity by direct beam counter |
| Absolute intensity scaling method | Comparison with scattering from pure $H_2O$ |
| Exposure time | Regions II-V: 40 sec<br>Regions I and VI: 60 sec<br>(Division into the regions is shown in Figure 1A) |
| Sample temperature (°C) | 25 |
| **(c) Software employed for SAS data reduction, analysis and interpretation** | |
| 1D-integration, averaging, data subtraction and Guinier analysis | RAW 2.2.1 |
| Calculation of ε from sequence | ProtParam: https://web.expasy.org/protparam/ |
| Calculation v values from chemical composition | Peptide Property Calculator:<br>http://biotools.nubic.northwestern.edu/proteincalc.html |
| Calculation of scattering length density (SLD) values from chemical composition | SLD calculator: http://www.ncnr.nist.gov/resources/activation/ |
| P(r) calculation | GNOM 5.0 from ATSAS 2.8.4 |
| Atomic structure modeling and fitting the data | CRYSOL; OLIGOMER |
| Molecular graphics | VMD 1.9.4a53 |

| **(d) Structural parameters** | | | | | | |
|---|---|---|---|---|---|---|
| Ranges | I | II | III | IV | V | VI |
| Border positions (ml) | 9.17 – 9.71 | 10.3 – 10.6 | 11.2 – 11.5 | 12.1 – 12.5 | 12.8 – 13.2 | 13.9 – 14.5 |

| | | | | | | |
|---|---|---|---|---|---|---|
| Guinier analysis | | | | | | |
| I(0) ($10^{-3}$ × $cm^{-1}$) | 0.255 ± 0.010 | 7.99 ± 0.12 | 7.02 ± 0.11 | 6.48 ± 0.06 | 5.20 ± 0.05 | 9.53 ± 0.03 |
| $R_g$ (Å) | 73 ± 3 | 60.6 ± 1.2 | 48.2 ± 1.2 | 40.6 ± 0.6 | 37.5 ± 0.7 | 28.17 ± 0.13 |
| q $R_g$ range | 0.91-1.30 | 0.44-1.33 | 0.67-1.23 | 0.46-1.25 | 0.36-1.22 | 0.22-1.3 |
| P(r) analysis | | | | | | |
| I(0) ($10^{-4}$ × $cm^{-1}$) | 2.67 ± 0.02 | 84 ± 2 | 73 ± 2 | 64.6 ± 0.6 | 52.2 ± 0.6 | 95.3 ± 0.3 |
| $R_g$ (Å) | 80.2 ± 1.9 | 71 ± 3 | 55 ± 3 | 41.2 ± 0.6 | 39.1 ± 0.8 | 28.5 ± 0.2 |
| $d_{max}$ (Å) | 275 | 295 | 230 | 140 | 140 | 110 |
| q-range ($Å^{-1}$) | 0.0125 – 0.1095 | 0.0072 – 0.1253 | 0.0146 – 0.1464 | 0.0114 – 0.1604 | 0.0097 – 0.2166 | 0.0079 – 0.2694 |
| $q_{min}$ $d_{max}$ / π | 1.09 | 0.676 | 1.07 | 0.508 | 0.432 | 0.277 |
| Total quality estimate (GNOM) | 0.633 | 0.584 | 0.594 | 0.638 | 0.655 | 0.592 |
| Volume ($V_P$) ($nm^3$) [1] | 770 | 350 | 190 | 130 | 100 | 75 |
| MW calculated from $V_P$ (kDa) [1] | 640 | 290 | 160 | 110 | 84 | 62 |
| MW calculated from $V_C$ (kDa) [1] | 340 | 280 | 150 | 104 | 56 | 57 |
| **(e) Atomistic modeling (range VI)** | | | | | | |
| Method | CRYSOL program from ATSAS 3.3.0 was used to approximate experimental data by curve calculated for crystal structure of dimer. | | | | | |
| q-range for fitting, $Å^{-1}$ | 0.0079 – 0.2 | | | | | |
| Contrast of the solvation shell (e $Å^{-3}$) | 0.1 | | | | | |
| $\chi^2$ value | 2.48 | | | | | |

| | | | | |
|---|---|---|---|---|
| **(f) Oligomer content analysis (ranges III-V)** | | | | |
| Method | | OLIGOMER program from ATSAS 3.3.0 was used to approximate experimental data; fit by crystallographic structures of dimer (derived from crystal structure using PyMOL) and tetramer and structure of octamer, generated using Chimera 1.14, based on crystal contacts. Parameters: maximum order of harmonics = 50; addition of a constant component was used. | | |
| Ranges | | III | IV | V |
| q-range for fitting, $Å^{-1}$ | | 0.0139 - 0.1464 | 0.0114 - 0.1604 | 0.0097 - 0.2167 |
| $\chi^2$ value | | 1.45 | 1.52 | 1.09 |
| Oligomer fractions | dimer | 0 [2] | 4 ± 5 | 12 ± 5 |
| | tetramer | 46.7 ± 1.7 | 76 ± 6 | 80 ± 7 |
| | octamer | 53.3 ± 1.2 | 19.7 ± 1.3 | 8 ± 3 |

[1] Relative errors of VP and MW values are estimated at 10%.

[2] Not accounted for due to its low fraction.

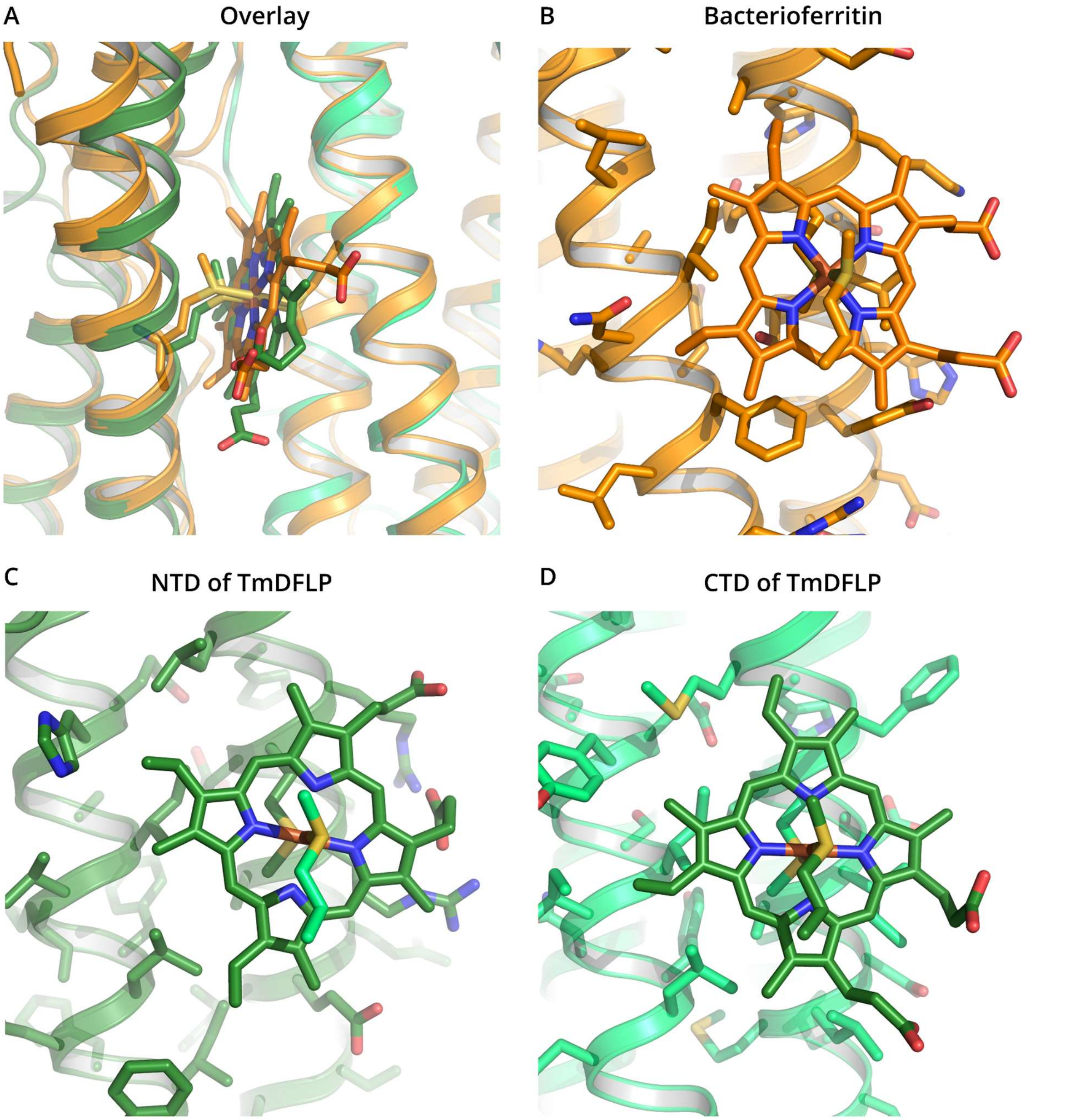


**Figure S1.** Comparison of putative heme binding site in TmDFLP (green, AlphaFold3 model) with the heme binding site of *Escherichia coli* bacterioferritin (orange, PDB ID 1BFR). A) Overlay of the two structural models. Backbone positions are similar between the proteins. Heme is symmetrically ligated by two methionine side chains. B) Heme binding site in bacterioferritin. Since the site is symmetric, only one side is shown. C and D) Putative heme-facing residues in NTD and CTD of TmDFLP.

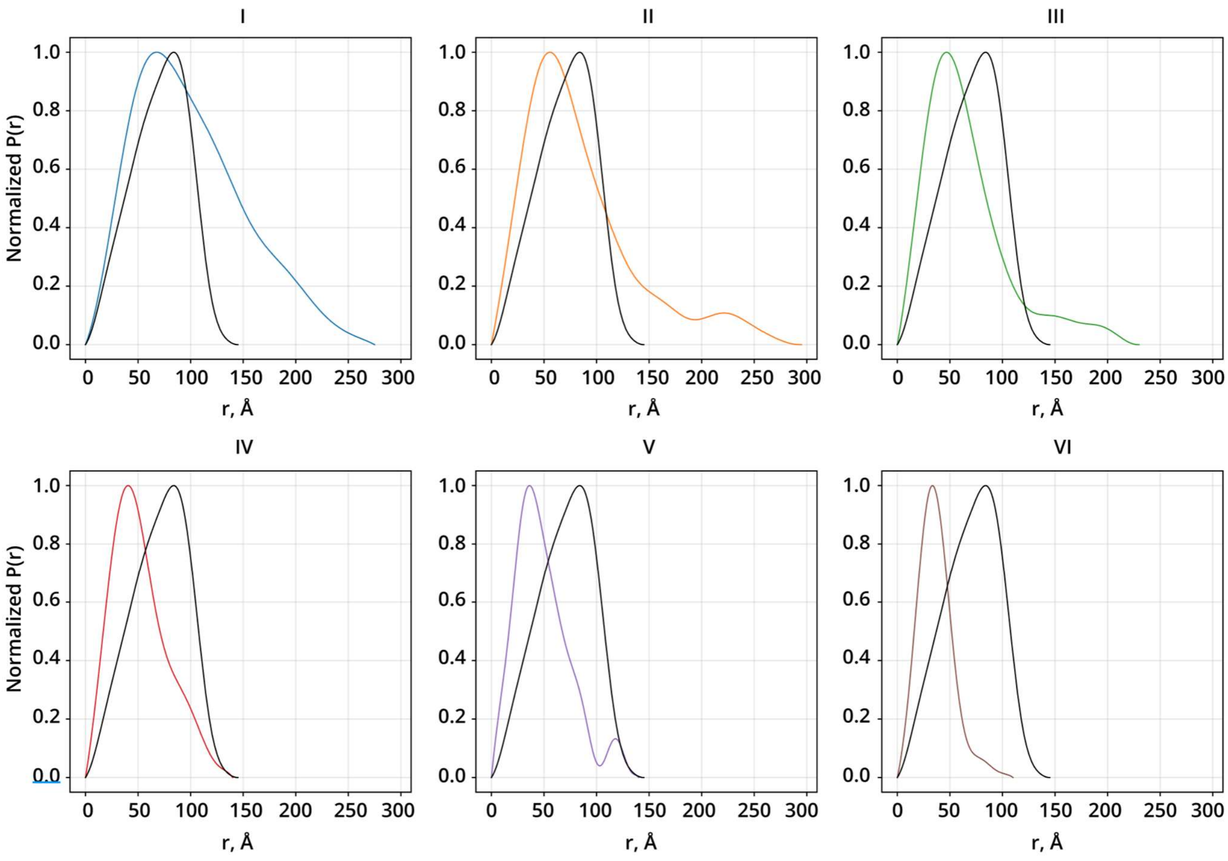


**Figure S2.** Comparison of normalized P(r) for ranges I-VI (colored) and *H. pylori* apoferritin curve (black) (SASBDB ID SASDTW4).

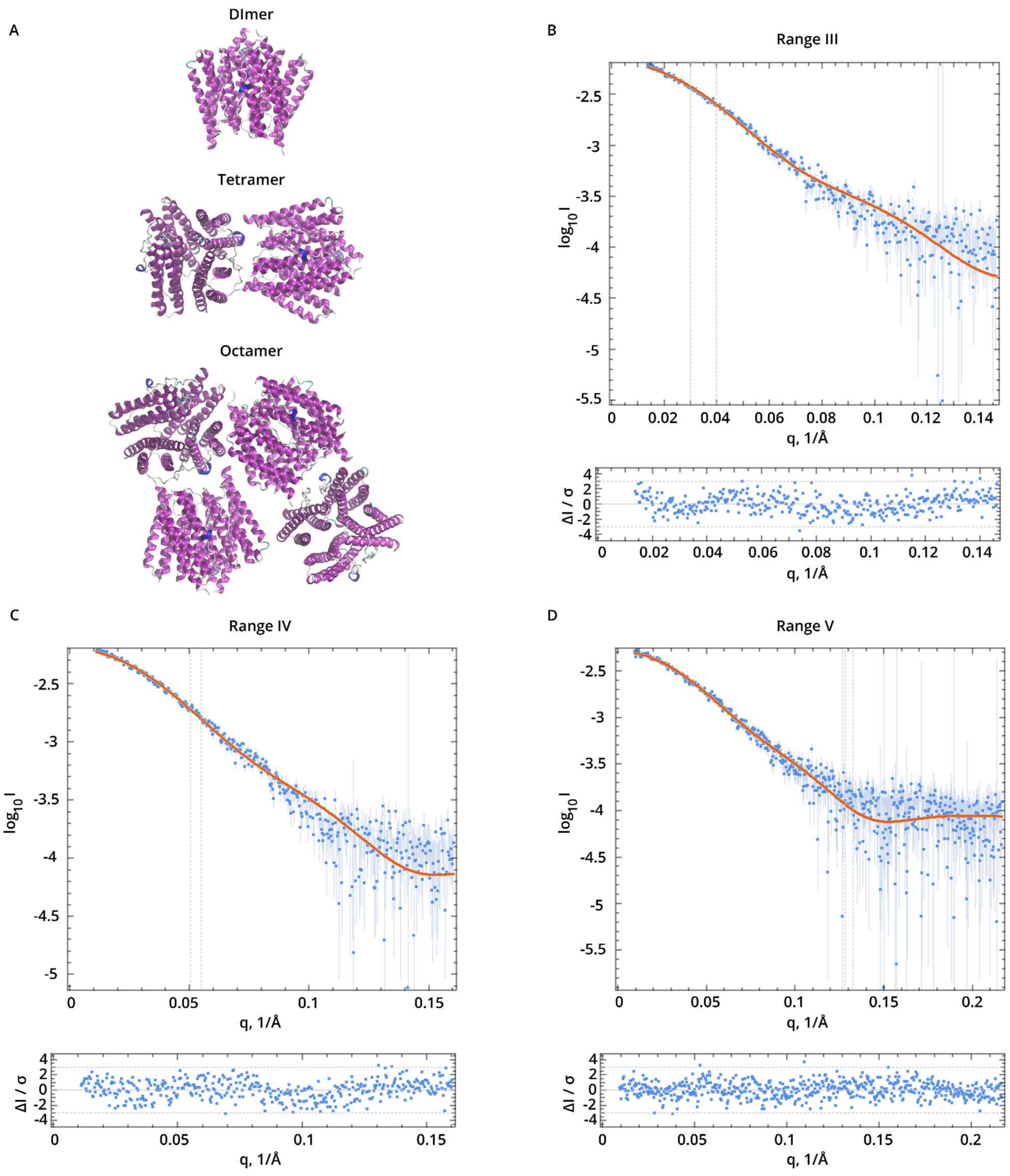


**Figure S3.** Oligomer content analysis for the SEC-SAXS ranges III-V (Figure 1A). A) Cartoon representations of the dimers, tetramers, and octamers of the TmDFLP generated based on crystal contacts. B, C, D) Fit by model curves in OLIGOMER for the ranges III, IV, and V, respectively. Relative residuals are shown at the bottom. Approximation results are presented in the Table S2(f).

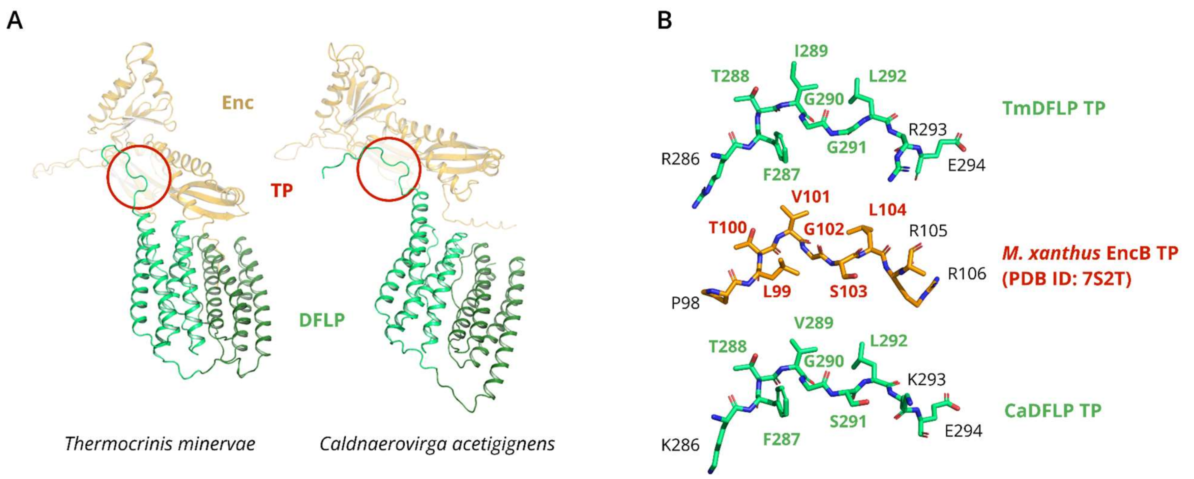


**Figure S4.** Tentative encapsulin-targeting peptides in TmDFLP and CaDFLP. A) Alphafold3 models of encapsulin-DFLP complexes for *T. minervae* and *C. acetigignens.* TPs marked with red circles. B) Comparison of TPs residues for TmDFLP, CaDFLP and *M. xanthus* EncB (PDB ID 7S2T).

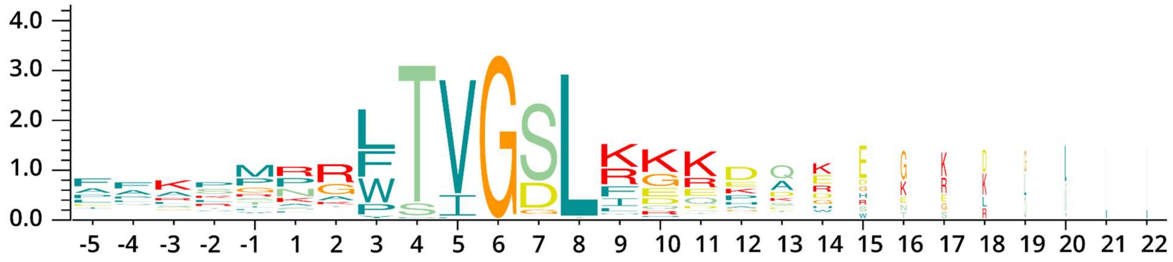


**Figure S5.** Extended weblogo of the C-termini of the encapsulin-targeted proteins in the frequency plot mode. Respective sequence alignment is provided in Figure S6.

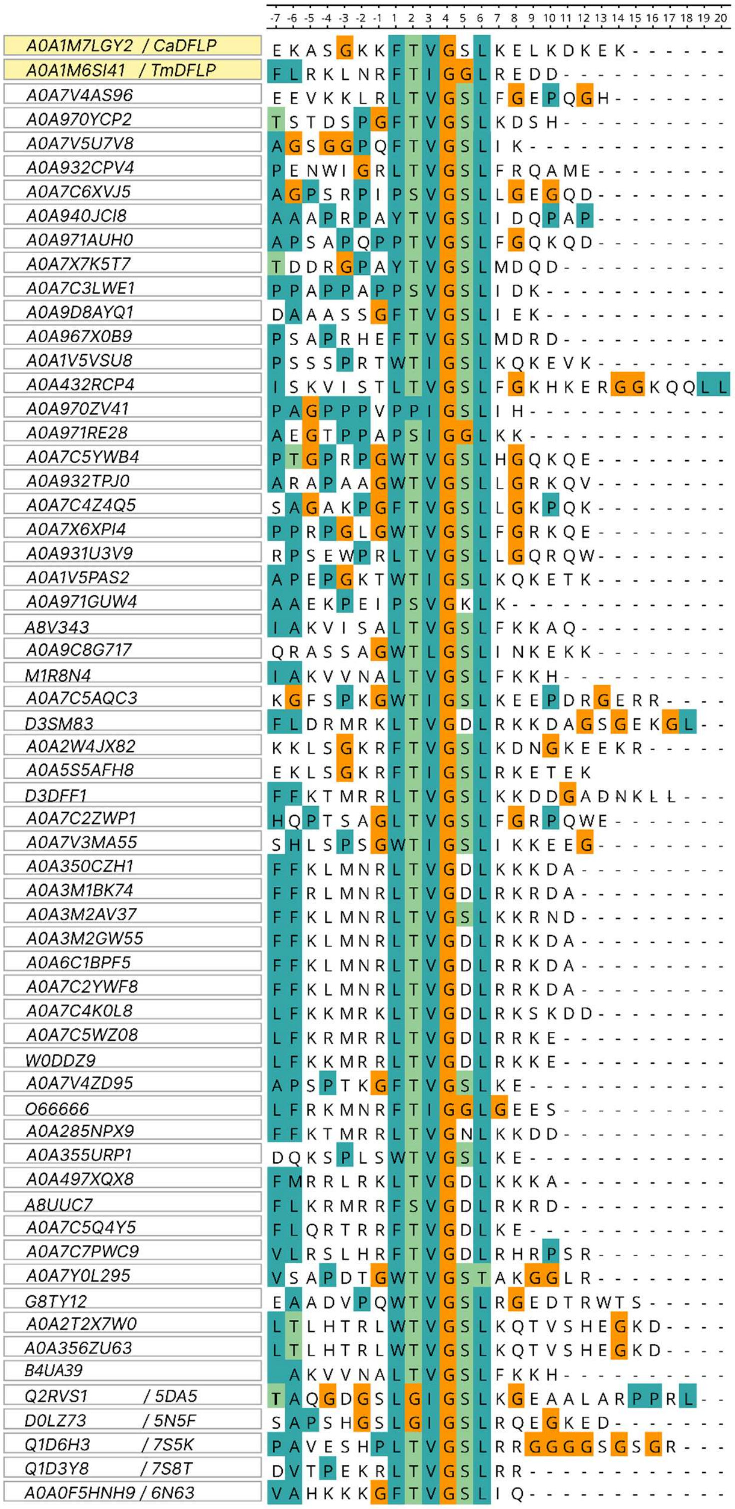


**Figure S6.** Sequence alignment of the putative encapsulin-targeting peptide regions of 56 DFLPs and 5 known encapsulin-interacting FLPs. PDB IDs are indicated where available.
